# Loss of Brain Structural qMRI Signatures in Aging

**DOI:** 10.1101/2025.09.18.677158

**Authors:** Niv Amos, Shir Filo, Tommy Kaplan, Aviv A Mezer

## Abstract

The human brain undergoes significant changes during aging, affecting its macrostructure and microstructure features and impacting its functionality. These aging-related changes are oftentimes linked to different neurodegenerative diseases. Previous studies using MRI and molecular analyses have proposed that aging in each individual brain leads to increased similarity between different brain regions, whereas on the population level, they show greater variability. We investigated these hypotheses using multi-parametric, microstructural, quantitative MRI (qMRI) approaches. Using seven qMRI parametric methods in data from young and older adults, we created unique microstructural signatures of 128 brain regions. Our results supported both hypotheses: First, that inter-regional microstructural similarity is stronger within older individuals; and second, that the variability in microstructural signatures across individuals is also greater in older than in younger adults. Our findings provide new insights into the brain’s microstructural changes during aging and demonstrate the potential of using multi-parameter qMRI techniques in neuroscience research.

## Introduction

As the brain ages, it undergoes notable structural changes that affect its function. Such changes have been documented in previous studies (Blinkouskaya et al., 2021; MacDonald and Pike, 2021) and include a reduction in overall brain volume, thinning of the cortical sheet, and alterations in white matter integrity. At the cellular level, aging is associated with a loss of neurons and synapses, along with changes in dendritic architecture and myelin integrity (West, 1993; Peters et al., 2008; Anderson and Rutledge, 1996; Bartzokis et al., 2010). These structural alterations may lead to decreased connectivity and communication between brain regions, potentially impairing cognitive functions (Damoiseaux, 2017). Understanding the structural processes underlying brain aging remains a longstanding objective in neuroscience research.

Various theories discuss the relationship between brain regions during the aging process. For example, Izgi et al. (2022) examined RNA expression patterns in rodents and proposed that aging is associated with a decline in the distinct organization of various body regions, including the cortex. This phenomenon has been referred to as the “identity loss” of those tissues. Nadig et al. (2021) suggested that cortical aging is a highly individualized process, meaning that as people get older, their brain structures become increasingly different from one another, making the aged cortex of one person less similar to that of other older adults of the same age. Fig. 1 illustrates these theories. While these observations have been shown in the cortex, they were not tested using MRI-based microstructural measurements.

**Figure 1:**
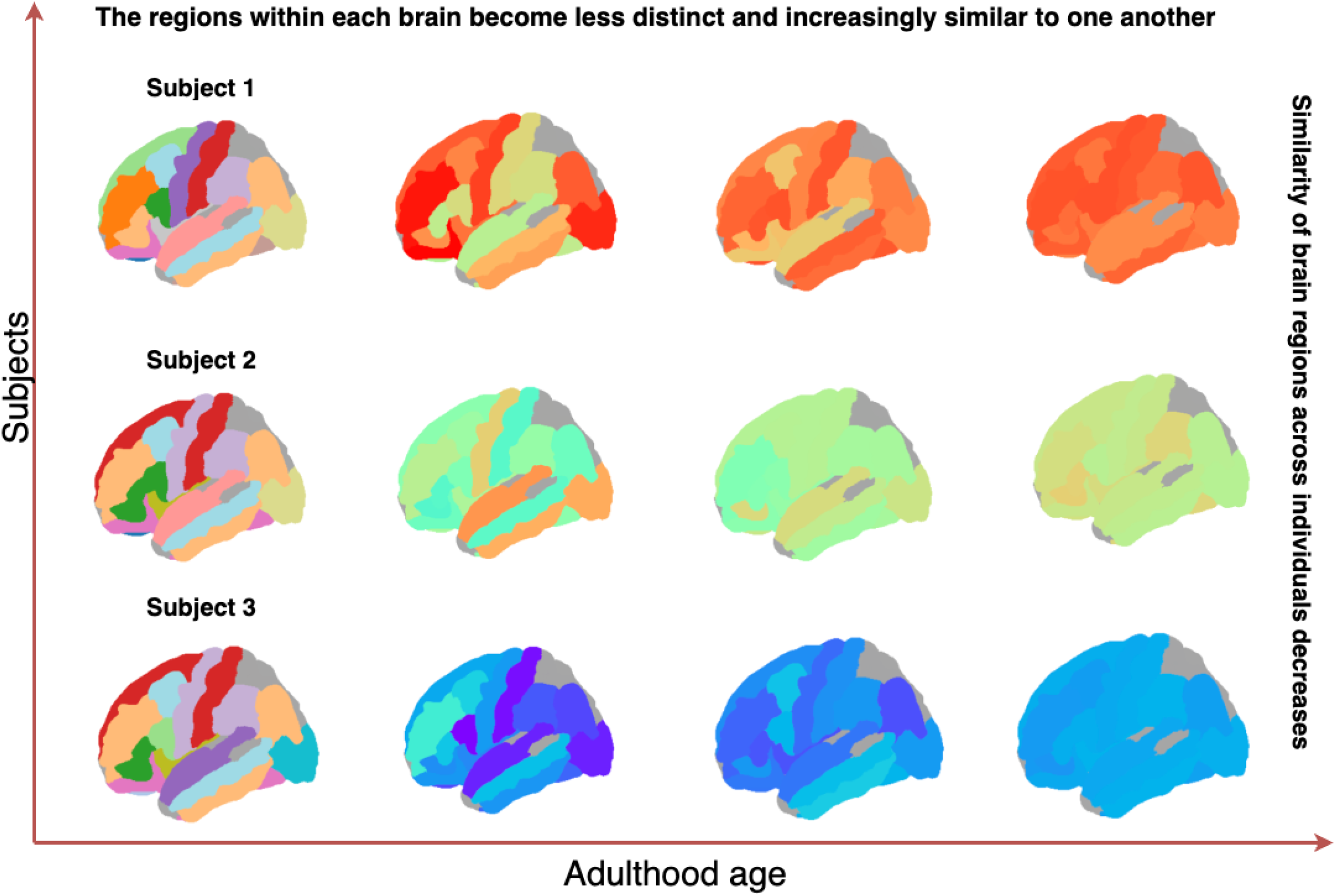
An illustration of our hypotheses. From left to right, the figure shows the progression of aging: initially, the brain regions exhibit distinct characteristics (represented by different colors). The rows represent three subjects’ brains, where these regions appear relatively similar at an early age. As aging progresses, the regions within each brain converge toward similar expression levels, becoming less distinct (with similar colors, representing similar values), while the similarity of these regions across different individuals decreases.

*In vivo* magnetic resonance imaging (MRI) serves as the primary tool for characterizing structural brain changes during aging, including alterations in brain volume and the thickness of various brain regions (Fjell and Walhovd, 2010; Lockhart and DeCarli, 2014; Sowell et al., 2004; Raz et al., 2005). In recent years, quantitative MRI (qMRI) and diffusion MRI (dMRI) have been introduced as methods to provide biophysically informed, microstructural-sensitive parametric measurements of the human brain during aging (Callaghan et al., 2014; Yeatman et al., 2014; Filo et al., 2019; Lebel et al., 2012; Chan et al., 2025). Interestingly, when comparing the aging contribution between macrostructural cortical thickness measurement and the qMRI R1, Erramuzpe et al. (2021) shows that each provided different and complementary aging information.

Recent studies have demonstrated the advantages of multiparametric approaches in brain imaging using multiple features for each region of the brain to serve as its unique signature, to assess regional similarity and covariance in individuals (Seidlitz et al., 2018; Sebenius et al., 2023, 2024). Paquola et al. (2019) extended these approaches to microstructural qMRI data (specifically, magnetization transfer (MT)) in order to characterize the myeloarchitecture of different cortical regions. Their findings revealed that myeloarchitectural properties can co-vary across the brain and that these properties provide a measure of regional similarity within an individual. Following such observations, the question is raised about how multiple qMRI microstructural parameters are interrelated and how they change relative to each other throughout the aging process.

Here, we investigate regional identity loss in the brain during aging using a multi-parametric qMRI approach, examining cortical and non-cortical regions in young and older adults. Specifically, we utilized seven neuroimaging parameters: three relaxation rates—longitudinal 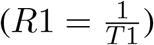, transverse 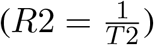, and effective transverse 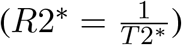; two macromolecular fraction parameters—the macromolecular and lipid volume fraction (MTV = 1 - water fraction) and the magnetization transfer (MT) saturation parameter; and two water diffusion characteristics—fractional anisotropy (FA) and mean diffusivity (MD).

Using these seven parameters, we examined two hypotheses: (1) whether microstructural signatures of different brain regions are more distinct in young adults but become more similar in older adults, and (2) whether inter-individual similarity in microstructural signatures decreases with age.

## Results

To test the relationship between the microstructure signature of the brain and age, we use a qMRI multiparameter vector, representing the microstructure signature for each selected brain region. Specifically, we tested two hypotheses. First, for a given brain, different regions’ microstructure signatures will be distinct in the young and become more similar with aging. Second, during aging, the microstructure signature similarity between individuals will be reduced.

### Constructing the multi-parametric vector

To test these hypotheses, we preprocessed our dataset through the following steps: We used 32 young and older adult subjects. For each of them, we used seven qMRI and dMRI maps. From each of these maps, we extracted data (voxel values) from 128 distinct brain regions of interest (ROI). The ROIs were selected from different areas, from both hemispheres, such as cortical, white-matter, and subcortical regions. The regions were defined using the automated cortical parcellation method (Desikan et al., 2006) as implemented in FreeSurfer (Fischl, 2012). Then, for each area (cortical, white-matter, and sub-cortical), we standardized the values within each map by calculating z-scores (based on values from the entire regions of this area) in order to address the differing scales across the maps. To further enhance data robustness and reduce computation time (as each map contains tens of thousands of voxels and values), we computed the median value for each region in each of the maps. Thus, for each subject, we constructed a 128 *×* 7 matrix where each row represents a given region’s seven-value microstructure signature. For a visual representation of the preprocessing steps, see Fig. 2

**Figure 2:**
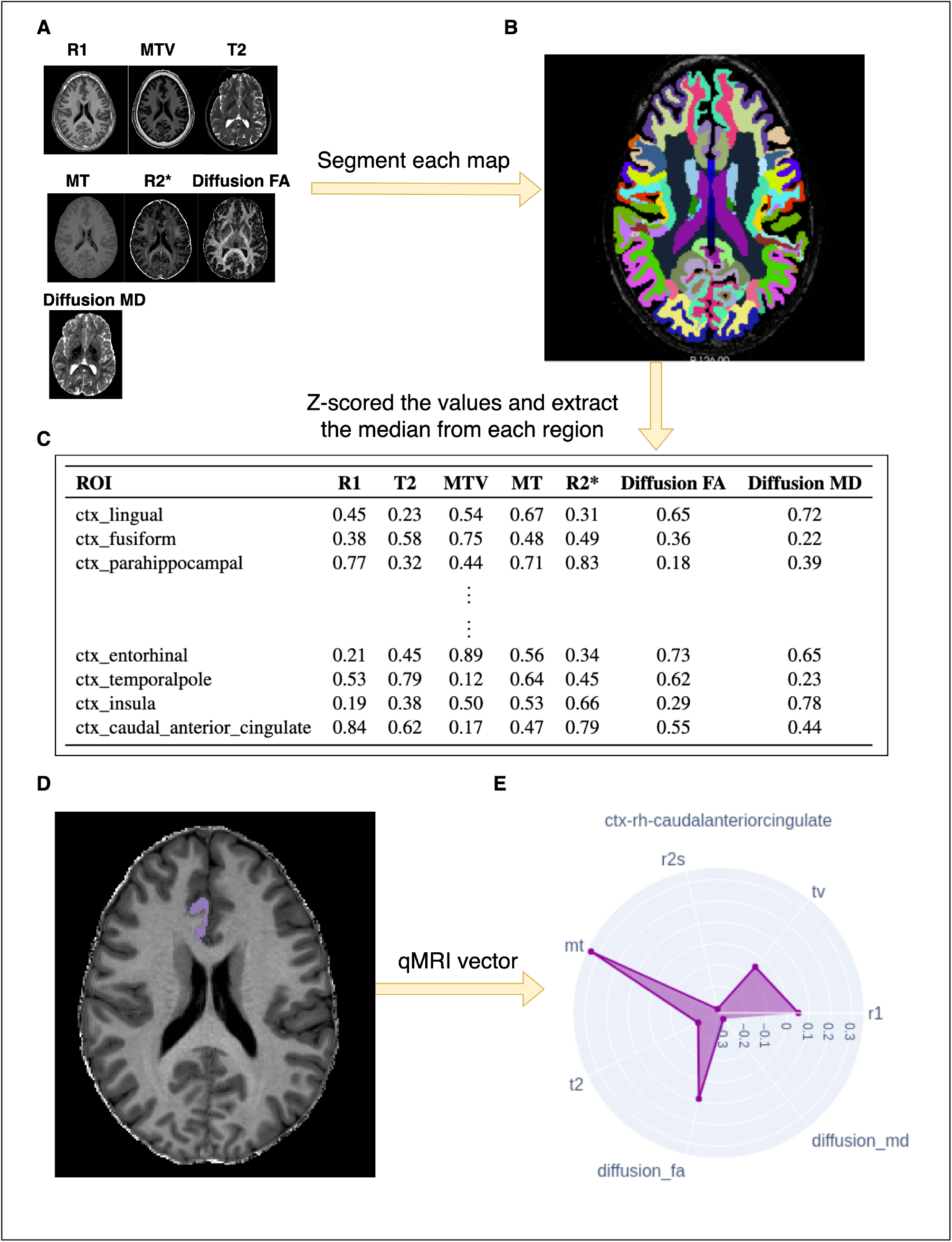
Illustration of our preprocessing method for a single subject. **A**. Visualization of the seven qMRI and dMRI maps; **B**. Example of a parcellated brain map; **C**. 128×7 matrix representing the subject’s brain regions (Z-scores and medians); **D**. An example of a cortical region; **E**. Our proposed microstructure signature, displayed using a polar plot, for the region in D.

### Testing the reliability of the multiparameter vector

We created a multi-parameter qMRI and dMRI vector as the microstructure signature of 128 brain regions in order to investigate the relationship between brain ROIs microstructure properties and age. We started by testing the vector’s robustness and reliability as a brain measure to identify ROIs and age.

First, we tested whether using our microstructure signatures—comprising different numbers of microstructural parameters—improves the accuracy of classifying ROIs as belonging to a young or old age group. For this, we used the XGBoost classification algorithm (Wu et al., 2021). The dataset consisted of ROIs (rows), each described by seven parameters. We trained one model at a time on the full dataset for each parameter set—either all seven parameters or specific subsets—using the same training/testing split.

The model trained with the full seven-parameter microstructure signature achieved the highest performance, with an average precision of 84%, outperforming models trained on reduced signatures (see Table 1). These results indicate that including more parameters in the microstructure signature improves classification accuracy.

**Table 1:** Binary classification evaluation matrix on young and old subjects using XGBoost, comparing the usage of a multi-parameter vector versus vectors with fewer parameters.

| Number of parameters | 1 | 2 | 3 | 4 | 5 | 6 | 7 |
| --- | --- | --- | --- | --- | --- | --- | --- |
| Precision ($\frac{TP}{TP+FP}$) | 60% | 65% | 70% | 75% | 79% | 82% | **84%** |
| Recall ($\frac{TP}{TP+FN}$) | 60% | 64% | 68% | 72% | 75% | 78% | **79%** |
| F1-Score<br>($2 \times \frac{\text{Precision} \times \text{Recall}}{\text{Precision} + \text{Recall}}$) | 60% | 64% | 68% | 72% | 75% | 78% | **78%** |

To assess the ROI information of the microstructure signature, we examined whether the similarities between signatures of homologous cortical brain regions are greater than non-homologous ones. In Fig. 3, we plot the mean cosine similarity across subjects and ROIs of homologous and non-homologous cortical brain regions. We observed that homologous regions, on average, exhibited significantly higher similarity (*p <* 0.001).

**Figure 3:**
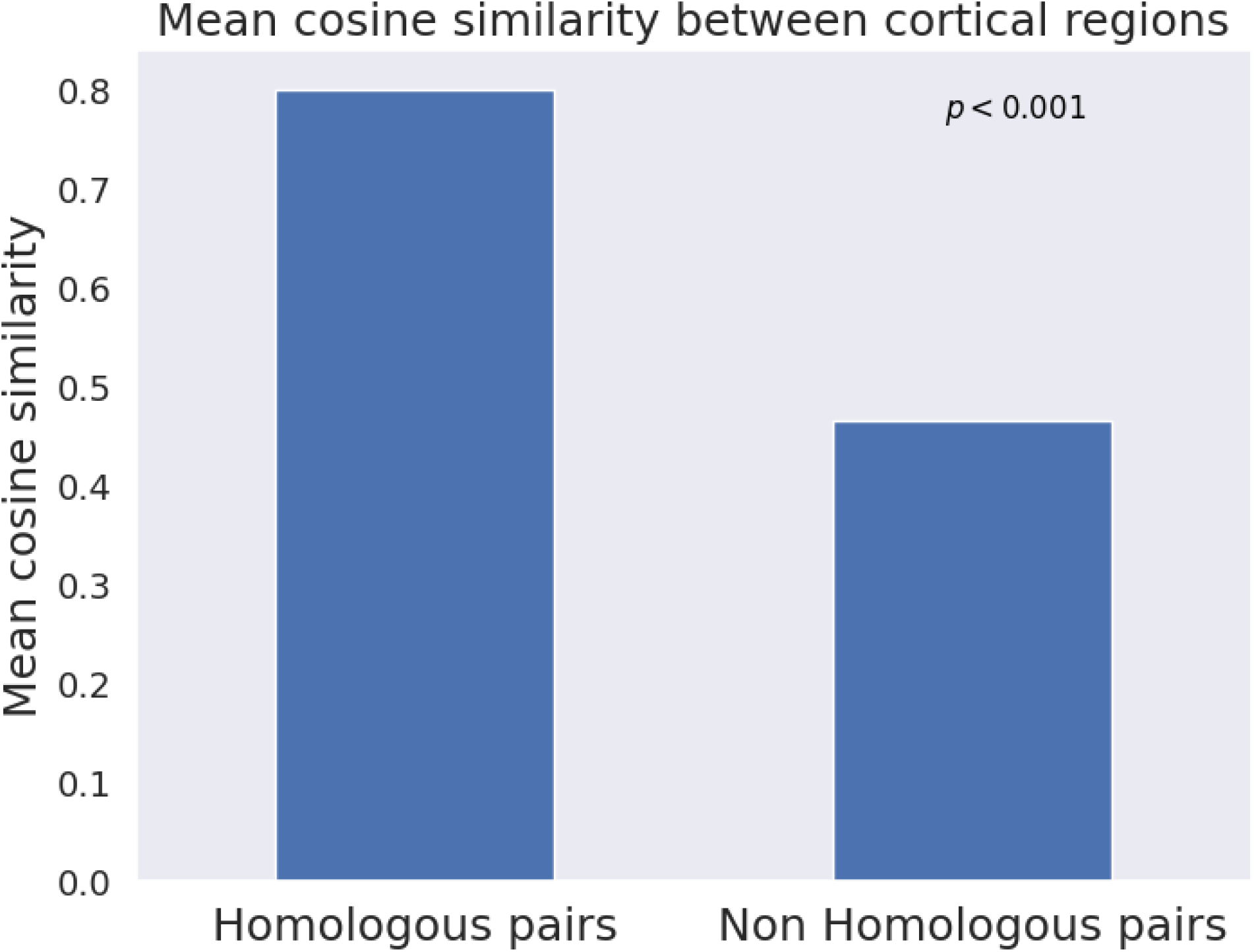
The mean cosine similarity between homologous and non-homologous cortical brain regions. On average, homologous regions have significantly higher similarity than non-homologous regions.

To further evaluate the ROI information, we used the t-SNE model to visualize how brain regions can be grouped according to their microstructure signature. In Fig. 4.A, we show that the model effectively differentiated between areas, with most of the cortical area in the center and the white matter regions surrounding it. The distribution of sub-cortical regions is of particular interest (Fig. 4.B) along the t-SNE model space. This representation suggests that each subcortical ROI has a different parametric signature. Also, some of the regions from the white matter area were also grouped together (Fig. 4.C).

**Figure 4:**
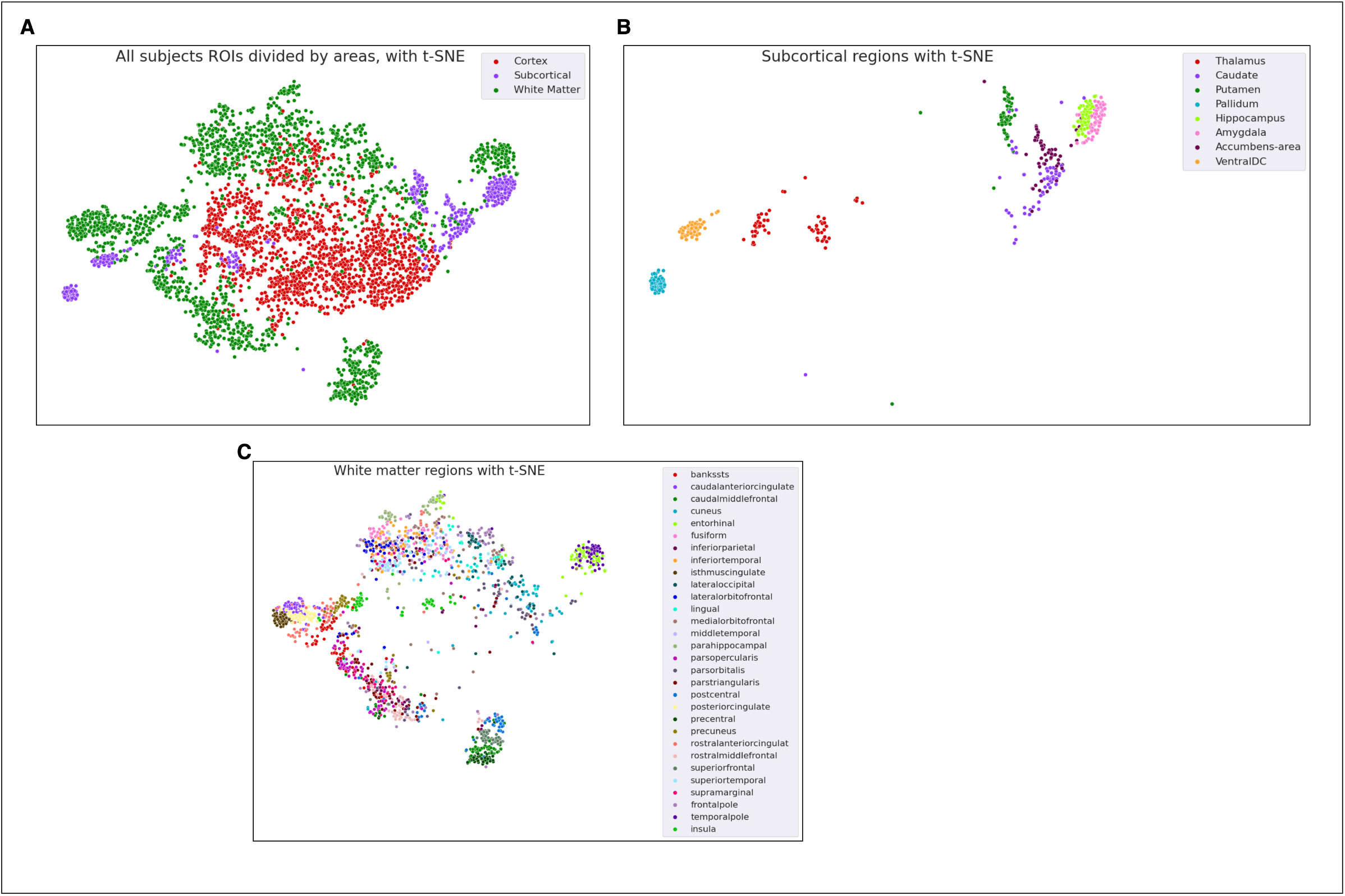
t-SNE visualization of ROI representations after dimensionality reduction. Each subject’s ROIs were projected into two dimensions using t-SNE. **A**. The resulting clusters are color-coded by their brain areas - cortical, white matter, and subcortical. **B**. A further breakdown of the subcortical regions from A. **C**. Some of the white matter regions were also clustered uniquely.

### Brain regions microstructure signature similarity during aging

Next, we tested our hypothesis that the microstructure signatures of brain regions are distinct from each other in young individuals and become more similar with age. We examined the inter-regional correlations among subjects and assessed the impact of aging.

We first calculated the Pearson correlation coefficients between the ROI vectors for each subject, generating a correlation matrix (Fig. 5A). We averaged these correlation coefficients across all subjects in each age group, resulting in two average correlation matrices (Fig. 5B). We then thresholded the negative correlations, similar to the work of Seidlitz et al. (2018). Finally, we collapsed each row from those matrices to derive the average correlation vector per group (Fig. 5C). This correlation vector represents the average correlation strength of each region with all other regions, thereby highlighting the brain regions with the strongest microstructural correlations.

**Figure 5:**
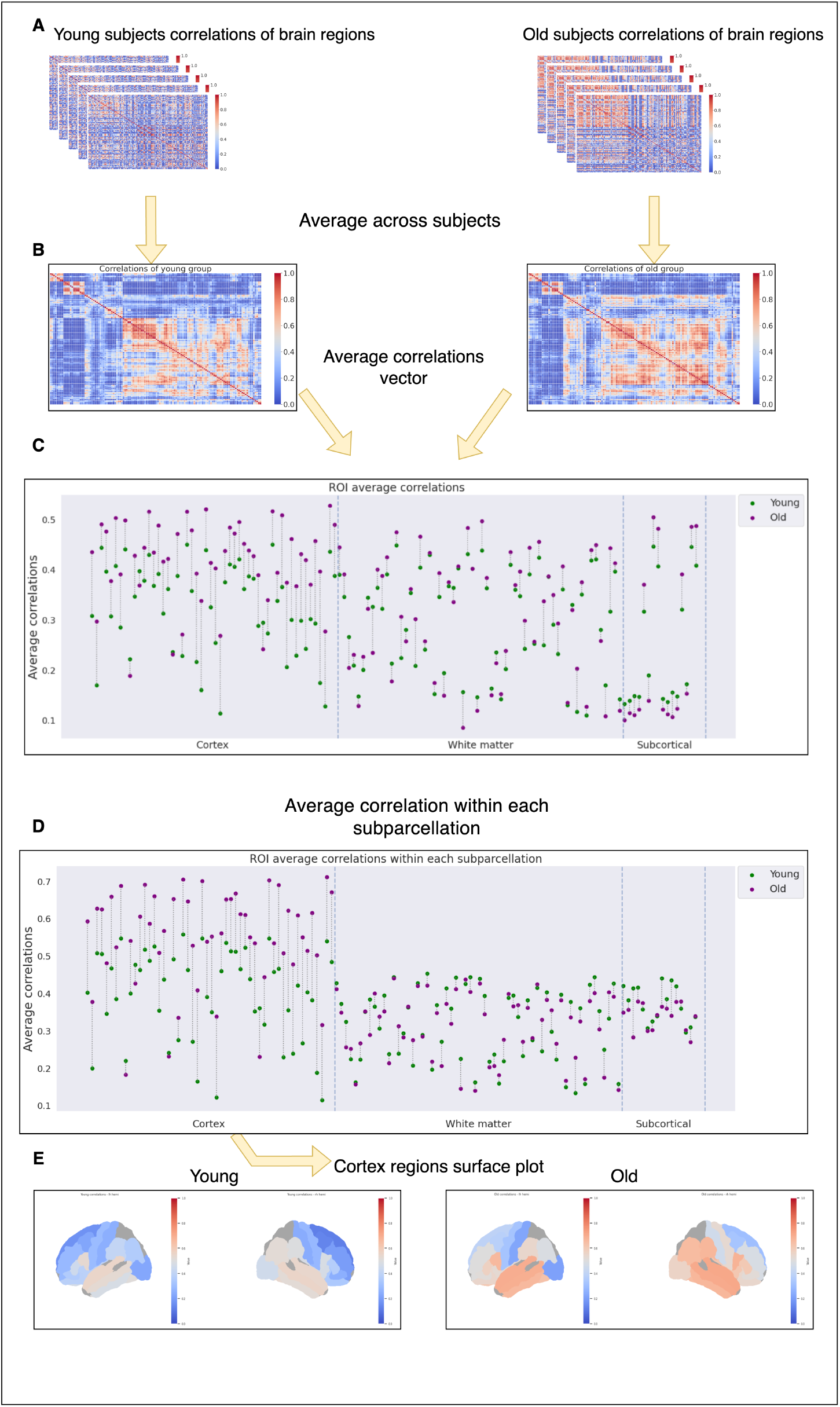
Overview of our ROIs correlations analysis. **A**. We constructed correlation matrices for each subject in both age groups, representing the correlations between all ROIs. **B**. We averaged these correlation matrices within each age group to produce a mean correlation matrix. **C**. For each ROI, we calculated the mean value of its correlation vector (each row in the average correlation matrix) to assess overall correlation strength. The results show that the microstructural signatures of brain regions in older subjects are generally more correlated than they are in younger subjects. **D**. We repeated this analysis for regions within each neuroanatomical subparcellation (cortex, white matter, sub-cortical gray matter) separately, revealing that cortical regions exhibit significantly higher correlations in older subjects than in younger ones. **E**. To visualize these findings, we mapped the average correlations for each cortical region onto a cortical surface, highlighting the increased connectivity in older subjects.

We performed an independent two-sample t-test to compare the overall average interregional correlations between the brain regions of older and younger subjects. The result of this test showed a significant difference between the groups (*p <* 0.001). In other words, older subjects demonstrated higher inter-regional correlations than younger subjects across almost all brain regions.

We then analyzed the average correlations within each area on its own subparcellation (cortex, white matter, gray subcortical) (Fig. 5D). When we compared the average correlations only between brain regions within each subparcellation, we found that the correlations in the cortex of older subjects differed significantly from those of younger subjects (*p <* 10^*−*^8), with nearly all regions showing higher correlations in the older group. For better visualization, in Fig. 5E we can see the cortical region’s correlations on a cortical surface.

Fig. 6 shows an example of the typical difference between two cortical regions (right caudal anterior cingulate and right pars triangularis) of young and older adults. Interestingly, we didn’t find similar differences between the groups for the white matter and subcortical regions.

**Figure 6:**
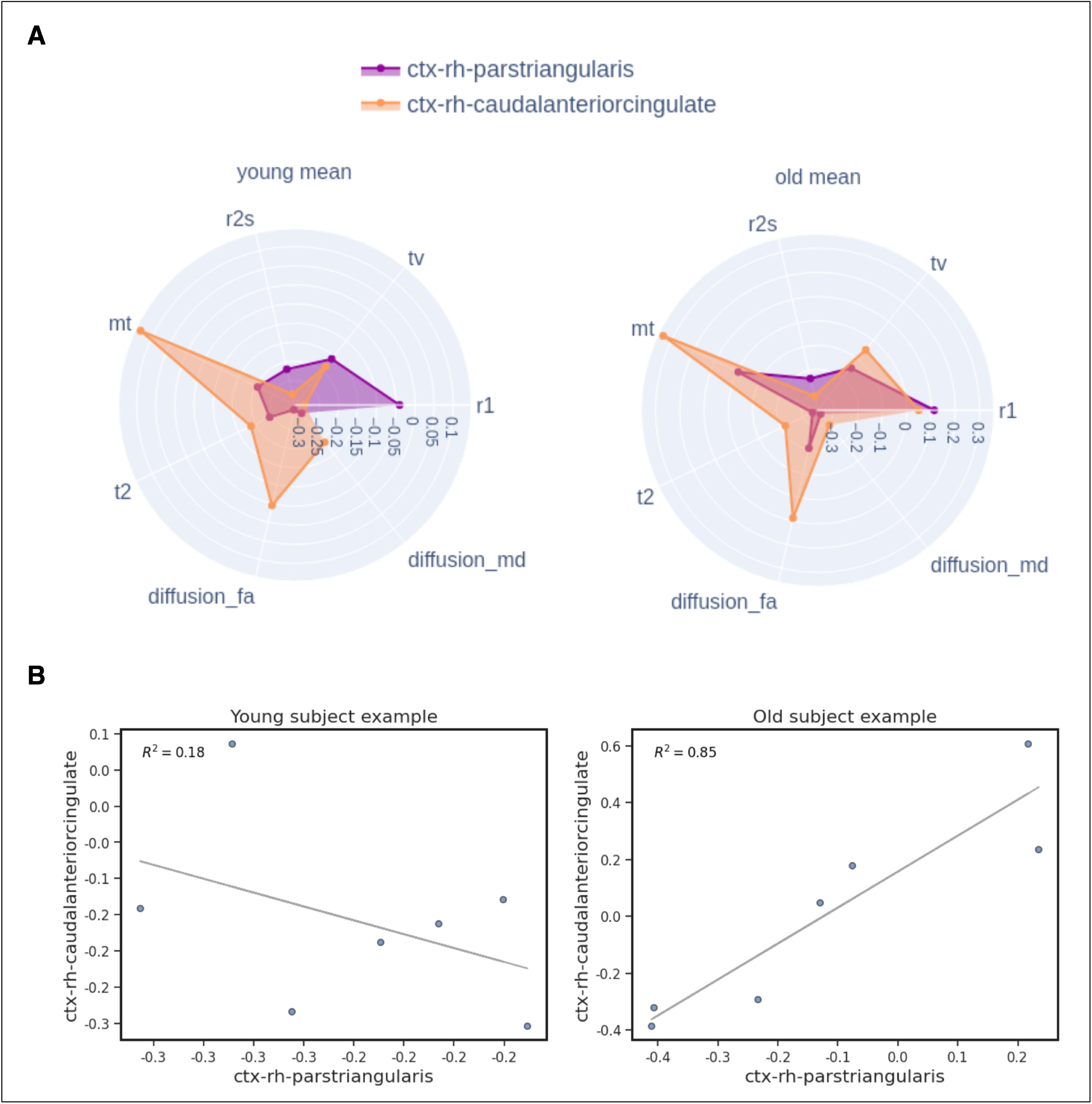
Comparison of two cortical regions’ correlation: right caudal anterior cingulate and right pars triangularis. **A**. Mean values from the median multi-parameter vector for both regions were calculated and visualized using a polar plot for each age group. The figure shows greater overlap in the older subjects, indicating increased similarity. **B**. A linear regression model was applied to the vectors from both regions for a random example subject from each age group. The model fit the vectors from these regions significantly better in older subjects, with an *R*^2^ value of 0.85 compared to 0.18 in younger subjects, indicating a stronger relationship between these regions in older individuals.

Finally, we applied a Bonferroni correction to our significance threshold to account for multiple comparisons across the different areas. Given three comparisons, we adjusted the alpha value from 0.01 to 0.003. The cortex p-value remained well below this threshold, indicating a significant effect that is unlikely to be a false positive. To validate our findings, we conducted a bootstrap analysis by resampling subjects with replacement over 1,000 iterations, which confirmed our findings.

### Microstructure signature similarity across individuals with aging

Our second hypothesis states that during aging, the microstructure signature similarity decreases across individuals. To assess this hypothesis, we calculated the standard deviation of each region’s microstructure signature values across all subjects in each age group, averaging these values for each parameter. Our analysis revealed that in most regions, older subjects exhibited greater variability (Fig. 7). Notably, the subcortical regions showed the largest difference in variability between the older and younger groups, indicating that these regions are more disrupted in older individuals.

**Figure 7:**
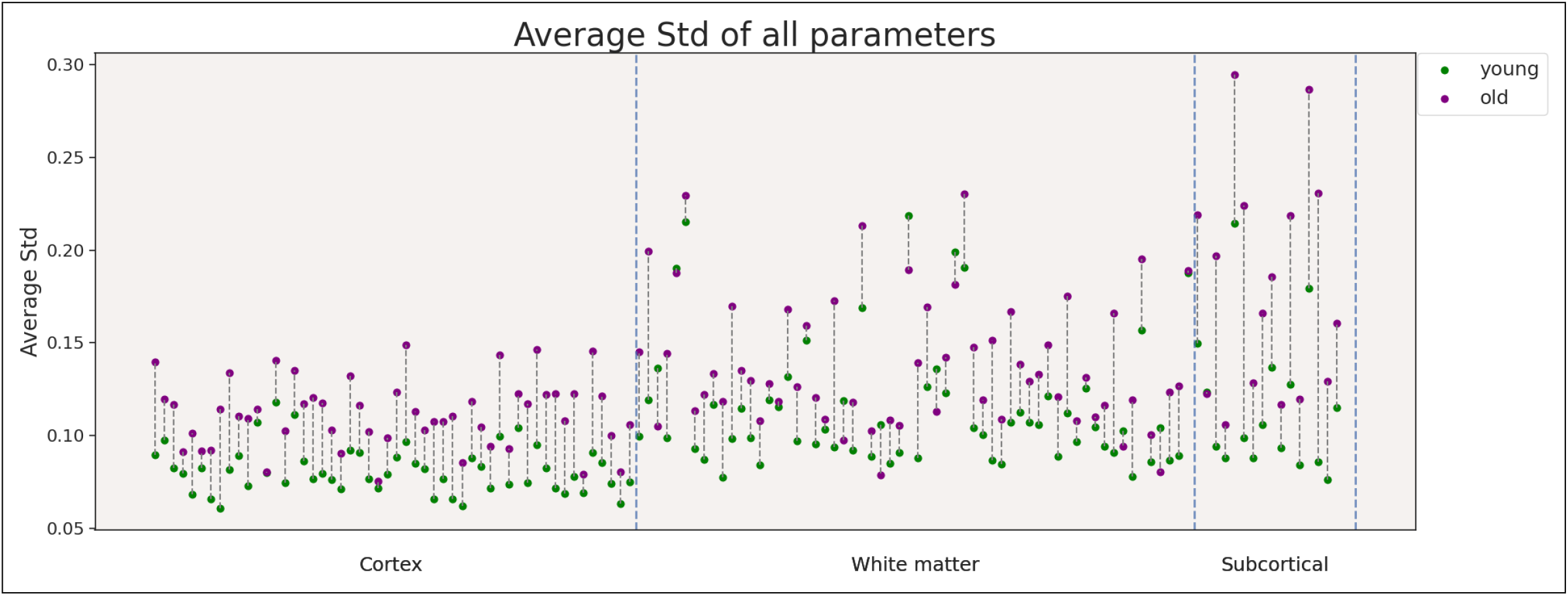
Variability of brain regions between age groups. The scatterplot compares the average standard deviations of the multi-parametric vectors for each brain region between young and old subjects. Each point represents a specific region, with greater variability observed in older subjects across most regions. Notably, subcortical regions showed the largest differences in variability, indicating increased disruption in these areas within the older group.

## Discussion

Our study utilized a qMRI and dMRI dataset to investigate the microstructural changes in the brain associated with aging. We utilized methods used for cortical macrostructural differences to analyze a multi-dimensional microstructural dataset. We first tested the stability of our approach to identify age and regional brain signatures. Next, we used these microstructure signatures to test two hypotheses regarding the similarities between brain regions within and between individuals. Our findings support the hypotheses, showing that as age increases, we observe an increase in both the correlation between the cortical regions and the variability across subjects.

In this work, we devised a new approach to assess brain regions’ microstructural signatures. Therefore, we assessed the approach’s usefulness and robustness. First, we showed that this approach allows a classification of the age group of each of the brain regions’ microstructural signatures. A multi-parameter approach can provide an improvement in precision of up to 20% compared to a single-parameter approach. Importantly, predicting the biological age of the brain and its sub-regions is a growing field of research with great clinical potential (Baecker et al., 2021; Lee et al., 2022; Aycheh et al., 2018; Erramuzpe et al., 2021). Hence, it will be interesting to further develop and test the potential benefits of our microstructural signature approach in this context.

Second, we applied the t-SNE visualization model to cluster regions according to their microstructural signature values. The t-SNE visualization shows that the data points of cortical regions are clustered together and surrounded by those of white-matter regions. Interestingly, each subcortical region has a very unique signature that distinguishes it from other brain regions. The unique microstructural signatures of subcortical regions echo earlier work by Bazin et al. (2020), which used the subcortical unique multi-parametric qMRI to segment those regions. Additionally, some white matter regions formed distinct clusters, indicating that this pattern is not occurring randomly but is instead driven by the intrinsic properties of their microstructural values.

Next, we used the microstructural signature across brain regions to test two aging-related hypotheses. We showed that microstructural signature similarities between different brain regions increase with age, especially in the cortical regions. This finding supports the hypothesis that aging is associated with a loss of regional specificity since the microstructural values of the aged brain tend to converge, resulting in a more homogeneous signature. This result agrees with previous studies in postmortem gene expression in the rodent cortex (Izgi et al., 2022). In that study, it was hypothesized that tissues converge during aging due to the loss of tissue identity. Our results suggest that a similar process also may affect the qMRI signature. Yet these observations may not be attributable solely to a loss of tissue identity: Other factors, such as general atrophy (Fjell et al., 2009) and shifts in key tissue components (e.g., myelin, iron, and water fraction), can play significant roles in these results (Bartzokis et al., 2010; Filo et al., 2019, 2023; Weiskopf et al., 2021).

Finally, we analyzed the standard deviation of the microstructural signature values across subjects in each age group. The results of this analysis support the hypothesis that the similarity of microstructural signatures of brain regions across different individuals decreases with age. Previous work by Nadig et al. (2021) showed that anatomical imbalance, which measures deviations from typical brain structure patterns, decreases during development but increases with aging, suggesting that brain morphology becomes more individualized with age. In agreement, we found greater variability in older subjects across most brain regions. This suggests that as individuals age, each brain changes uniquely. One can posit several possible sources for such differences between individuals, including genetics (Kremen et al., 2010; Izgi et al., 2022), lifestyle factors (Erickson et al., 2011; Valls-Pedret et al., 2015), environmental exposures (Kremen et al., 2010), and medical history (Wrigglesworth et al., 2023; Muller et al., 2014). Interestingly, the greatest variability across subjects was found in the subcortical regions. This may highlight an increased vulnerability of these regions to age-related changes or the MRI parameters’ sensitivity to changes in those regions. Indeed, subcortical regions are known to be involved in neurodegenerative diseases (Power and Looi, 2015; Rao et al., 2022; Redgrave et al., 2010). Furthermore, those regions are rich in paramagnetic compounds such as iron, which undergo changes in the aging process (Filo et al., 2023; Chan et al., 2025); which also may contribute to our observation.

Our results may depend on the set of qMRI and dMRI parameters that we collected for the data. Each parameter represents a contribution of multiple biological sources, at least some of which are likely to change with age (Weiskopf et al., 2021). Therefore, a larger dataset (e.g., (Jansen et al., 2024; Slater et al., 2019)) may enable the optimization of such parameter sets, potentially leading to different parameter sets being optimized for different scientific questions.

Our analysis is likely reflective of the methods that we selected to segment the brain regions. Previous studies showed that qMRI values can separate between brain regions (Schurr et al., 2018, 2019, 2020). Hence, different segmentation approaches (Yaakub et al., 2020) may yield different outcomes. Furthermore, qMRI values are known to vary across brain ROIs (Drori et al., 2022; McColgan et al., 2021); indeed, an earlier work by Paquola et al. (2019) showed that multiple samples across the cortex can be used to generate similar covariance matrices that change with age.

To conclude, our study provides evidence for increased inter-regional similarity within individuals (particularly in cortical areas) and increased variability across individuals in the aging brain. Our method utilizes multi-dimensional normalization, which has been shown to be useful for various tasks such as clustering and classification. These findings contribute to our understanding of normal brain aging and demonstrate the potential of multi-parameter qMRI approaches for investigating brain microstructure. Future research building on this work could significantly advance our knowledge of brain aging.

## Methods

### Overview

We used qMRI data to test our hypotheses concerning similarities in brain regions during aging. We used the human *in vivo* MRI dataset collected by Filo et al. (2019). All of our analyses were conducted using the Python programming language (https://github.com/python).

### Subjects

From the dataset, we selected 17 young adults (aged 26.4 *±* 2.4 years, 9 females, 8 males) and 15 older adults (aged 66.1*±* 5 years, 4 females, 11 males). For each subject, we used seven qMRI maps: R1 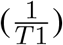, T2, MTV, MT, R2* 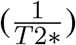, Diffusion FA, and Diffusion MD. All selected subjects are considered healthy.

### MRI acquisition

As described in Filo et al. (2019), data was collected on a 3 T Siemens MAGNETOM Skyra scanner equipped with a 32-channel head receive-only coil at the ELSC neuroimaging unit at the Hebrew University.

For quantitative R1, R2*, and MTV mapping, 3D Spoiled Gradient Echo (SPGR) images were acquired using different flip angles of 4°, 10°, 20°, and 30°. Each image had five echoes spaced with echo times ranging from 3.34 ms to 14.02 ms, and a repetition time of 19 ms. For T2 mapping, multi-SE images were obtained with ten spin echoes with echo times ranging from 12 ms to 120 ms, and a repetition time of 4.21s. MTsat mapping used SPGR images with an additional MT pulse, a flip angle of 10°, and echo and repetition times of 3.34 ms and 27 ms, respectively. Whole-brain DTI was performed using a diffusion-weighted spin-echo EPI sequence at 1.5 mm resolution, 64 directions, and a diffusion weighting of 2000 *s/mm*^2^, with echo time and a repetition time of 95.80 ms and 6000 ms, respectively, a gradient strength of 45 *mT/m*, a diffusion encoding duration of 32.25 ms, and a diffusion time of 52.02 ms.

### Estimation of qMRI parameters

Whole-brain MTV and R1 maps were computed using the mrQ software (Mezer et al., 2013). T2 maps were computed by implementing the echo-modulation curve (EMC) algorithm (Ben-Eliezer et al., 2015). MTsat maps were computed as described in (Helms et al., 2008). Diffusion analysis was done using the FDT toolbox in FSL (Smith et al., 2004). The R2* was estimated by using SPGR scans with multiple echoes. Fitting was done through the MPM toolbox (Weiskopf et al., 2013).

### Brain segmentation

Whole-brain segmentation was computed automatically using the FreeSurfer segmentation (Fischl, 2012). For subjects who had an MPRAGE scan, we used it as a reference. For the other subjects, the MP2RAGE scan was used as a reference. These anatomical images were registered to the MTV space prior to the segmentation process, using a rigid-body alignment. Sub-cortical gray-matter structures were segmented with FSL’s FIRST (Patenaude et al., 2011).

### Selected brain regions

We used multiple cortical, subcortical, and white matter regions in our analysis as listed in: Table 2, Table 3, Table 4. The labels correspond to the Freesurfer labels. Each brain region we used has 6,400 voxels on average.

**Table 2:** Cortical ROIs labels.

| Cortical ROIs |  |  |
| --- | --- | --- |
| ctx-lh-caudalanteriorcingulate | ctx-lh-caudalmiddlefrontal | ctx-lh-cuneus |
| ctx-lh-fusiform | ctx-lh-inferiorparietal | ctx-lh-inferiortemporal |
| ctx-lh-isthmuscingulate | ctx-lh-lateraloccipital | ctx-lh-lateralorbitofrontal |
| ctx-lh-lingual | ctx-lh-medialorbitofrontal | ctx-lh-middletemporal |
| ctx-lh-parahippocampal | ctx-lh-parsopercularis | ctx-lh-parsorbitalis |
| ctx-lh-parstriangularis | ctx-lh-postcentral | ctx-lh-posteriorcingulate |
| ctx-lh-precentral | ctx-lh-precuneus | ctx-lh-rostralanteriorcingulate |
| ctx-lh-rostralmiddlefrontal | ctx-lh-superiorfrontal | ctx-lh-superiortemporal |
| ctx-lh-supramarginal | ctx-lh-insula | ctx-rh-caudalanteriorcingulate |
| ctx-rh-caudalmiddlefrontal | ctx-rh-cuneus | ctx-rh-fusiform |
| ctx-rh-inferiorparietal | ctx-rh-inferiortemporal | ctx-rh-isthmuscingulate |
| ctx-rh-lateraloccipital | ctx-rh-lateralorbitofrontal | ctx-rh-lingual |
| ctx-rh-medialorbitofrontal | ctx-rh-middletemporal | ctx-rh-parahippocampal |
| ctx-rh-parsopercularis | ctx-rh-parsorbitalis | ctx-rh-parstriangularis |
| ctx-rh-postcentral | ctx-rh-posteriorcingulate | ctx-rh-precentral |
| ctx-rh-precuneus | ctx-rh-rostralanteriorcingulate | ctx-rh-rostralmiddlefrontal |
| ctx-rh-superiorfrontal | ctx-rh-superiortemporal | ctx-rh-supramarginal |
| ctx-rh-insula |  |  |

**Table 3:** Subcortical ROIs labels.

| Subcortical ROIs |  |  |
| --- | --- | --- |
| Left-Thalamus | Left-Caudate | Left-Putamen |
| Left-Pallidum | Left-Hippocampus | Left-Amygdala |
| Left-Accumbens-area | Left-VentralDC | Right-Thalamus |
| Right-Caudate | Right-Putamen | Right-Pallidum |
| Right-Hippocampus | Right-Amygdala | Right-Accumbens-area |
| Right-VentralDC |  |  |

**Table 4:** White Matter ROIs labels.

| White Matter ROIs |  |  |
| --- | --- | --- |
| wm-lh-bankssts | wm-lh-caudalanteriorcingulate | wm-lh-caudalmiddlefrontal |
| wm-lh-cuneus | wm-lh-entorhinal | wm-lh-fusiform |
| wm-lh-inferiorparietal | wm-lh-inferiortemporal | wm-lh-isthmuscingulate |
| wm-lh-lateraloccipital | wm-lh-lateralorbitofrontal | wm-lh-lingual |
| wm-lh-medialorbitofrontal | wm-lh-middletemporal | wm-lh-parahippocampal |
| wm-lh-parsopercularis | wm-lh-parsorbitalis | wm-lh-parstriangularis |
| wm-lh-postcentral | wm-lh-posteriorcingulate | wm-lh-precentral |
| wm-lh-precuneus | wm-lh-rostralanteriorcingulate | wm-lh-rostralmiddlefrontal |
| wm-lh-superiorfrontal | wm-lh-superiortemporal | wm-lh-supramarginal |
| wm-lh-frontalpole | wm-lh-temporalpole | wm-lh-insula |
| wm-rh-bankssts | wm-rh-caudalanteriorcingulate | wm-rh-caudalmiddlefrontal |
| wm-rh-cuneus | wm-rh-entorhinal | wm-rh-fusiform |
| wm-rh-inferiorparietal | wm-rh-inferiortemporal | wm-rh-isthmuscingulate |
| wm-rh-lateraloccipital | wm-rh-lateralorbitofrontal | wm-rh-lingual |
| wm-rh-medialorbitofrontal | wm-rh-middletemporal | wm-rh-parahippocampal |
| wm-rh-parsopercularis | wm-rh-parsorbitalis | wm-rh-parstriangularis |
| wm-rh-postcentral | wm-rh-posteriorcingulate | wm-rh-precentral |
| wm-rh-precuneus | wm-rh-rostralanteriorcingulate | wm-rh-rostralmiddlefrontal |
| wm-rh-superiorfrontal | wm-rh-superiortemporal | wm-rh-supramarginal |
| wm-rh-frontalpole | wm-rh-temporalpole | wm-rh-insula |

### Dataset construction

To extract the 3D brain qMRI maps and segmentation maps for each subject, we used nibabel package (https://github.com/nipy/nibabel). The regions were divided to fit the segmentation maps using NumPy (Harris et al., 2020) methods.

We standardized voxel values within the brain regions using the z-score method 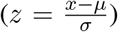 implemented in the SciPy library Virtanen et al. (2020). Initially, we combined all areas (cortical, white matter, and subcortical) for standardization, but this resulted in a non-normal distribution that persisted even after standardization. To address this, we standardized each area separately, as they each followed a normal distribution independently, leading to a normalized distribution for the combined regions.

The median values were calculated using NumPy (Harris et al., 2020), and the final resulting data matrix of each subject was stored in a Pandas dataFrame (McKinney and others, 2010).

### Outliers handling

Originally, the dataset contained 37 subjects, each with 140 brain regions. We removed subjects identified as noisy outliers using an interquartile range (IQR) method. For each parameter, we calculated the first quartile (Q1) and the third quartile (Q3) values. Outlier subjects were identified as those whose data points fall (on average) below Q1 minus a threshold (multiplied by the IQR), or above Q3 plus the same threshold. We applied the same method for the brain regions, to remove noisy regions. We ended up with 32 subjects, each with 128 brain regions.

### Correlations analysis

First, we used hierarchical clustering as a baseline for determining the order of the ROIs in the correlation matrices. This was based on the average linkage method, calculated as:

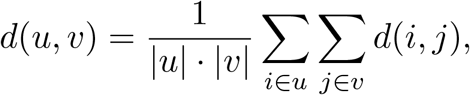

where the distance metric *d*(*i, j*) represents the cosine distance between points *I* and *j*. Cosine distance is defined as:

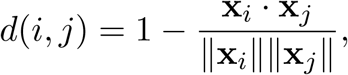

where **x**_*i*_ and **x**_*j*_ are the vectors representing the ROIs. This distance metric was used to capture the similarity between different ROIs based on their angular separation, ensuring the clustering reflects both magnitude and directional information in the data. This approach was implemented using the Scipy library Virtanen et al. (2020).

Then, we calculated the Pearson correlation coefficientbetween each two ROIs vectors, yielding a 128 *×* 128 correlation matrix. Each correlation was computed using the built-in pandas dataFrame (McKinney and others, 2010) correlation method.

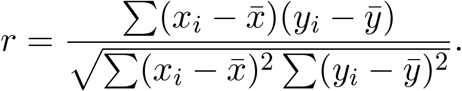

### Statistical significance

Through our analyses, we aimed to determine whether there were significant differences between the findings of the young and old groups. To achieve this, we used an independent two-sample t-test.

### Dimensionality reduction using t-SNE

For dimensionality reduction, we employed a t-distributed stochastic neighbor embedding (t-SNE) model, which we implemented by scikit-learn (Pedregosa et al., 2011). Before applying t-SNE, we performed a principal component analysis (PCA) to reduce the dimensionality while retaining 95% of the variance. This step was essential, as t-SNE tends to work better on lower-dimensional data (since it can be computationally expensive and sensitive to noise in high-dimensional spaces). We used the default hyperparameters for the t-SNE model, with the exceptions of setting the number of components to 2 (for 2-D visualization) and adjusting the perplexity to 100, which was selected after testing several values.

### XGboost model for Binary Classification

In order to perform ROI binary classification of age group, we used the default XGboost classifier (Chen and Guestrin, 2016). We trained each model using this classifier’s built-in training method. We split the dataset into 80% for training and 20% for testing.

### Bootstrap validation

To validate our findings, we conducted a bootstrap analysis by resampling subjects with replacement over 1,000 iterations. For each resampled dataset, we calculated the mean value of each region’s correlation vector for both the young and old groups. This resulted in 1,000 lists of mean correlation values from each brain region, for each group. We then averaged these lists across all iterations for both groups. For each brain region, we calculated the 95% confidence intervals of the mean correlation values based on these bootstrap samples.

## Data availability

The dataset analyzed during the current study is available from the corresponding author upon reasonable request.

## Code availability

The code for all of our analysis is available at: https://github.com/MezerLab/qMRI-age-similarities.

## Author Contributions

N.A., S.F., and A.A.M. conceptualized this research. N.A. developed and carried out all analyses under the guidance of T.M. and A.A.M. N.A. wrote the manuscript, with all authors contributing to revisions. S.F. and A.A.M. were responsible for data collection.

## Acknowledgements

This work was supported by the Israel Science Foundation (grants 1250/18, 259/23), the Center for Interdisciplinary Data Science Research, by the Ministry of Innovation, Science & Technology, Israel (grant 0005421) to TK and NA, and the Israel Science Foundation (grant no. 1169/20 and 2449/24) given to AAM and NA.

